# Hello From the Other Side: Robust Contralateral Interference in Tactile Detection

**DOI:** 10.1101/2020.07.21.213751

**Authors:** Flor Kusnir, Slav Pesin, Ayelet N. Landau

**Affiliations:** Departments of Psychology and Cognitive Science, The Hebrew University of Jerusalem, Jerusalem

**Author notes:** **Correspondence:** Ayelet N. Landau, Departments of Psychology and Cognitive Science, The Hebrew University of Jerusalem, Jerusalem.

**Keywords:** Tactile detection, interference, somatosensation, change detection, psychophysics

## Abstract

Our sense of touch is unique in that our tactile receptors are spread across our body surface and constantly receive different inputs at the same time. These inputs vary in relevance according to our current goals, but there is little research on how simultaneous stimulation to different body sites affects the perception of touch. In this series of studies, we characterised how irrelevant tactile sensations across the body-midline affect tactile detection in a constantly-attended body site. Participants had to detect a target on their dominant index finger, while receiving irrelevant stimulation to another body site (homologous and non-homologous fingers, and the contralateral ankle). We document robust interference effects on all measured body-sites. Its impact on detection-performance was unaffected by body posture, exacerbated by the intensity of the irrelevant stimulation, and ameliorated by embedding a target-like signal in the irrelevant stimulation. In addition, we generalise our findings beyond the target stimulus (i.e., a vibration intensity decrement) and report similar effects when employing a target-increment. In light of our findings, we propose that tactile inputs may be pooled together early in the hierarchy of somatosensory processing, resulting in an integrated percept. The rules for integration across body sides are likely not described by a simple summation, but rather may be governed by more complex interactions between fingers and according to the corresponding perceived, as well as actual, intensities of the stimulation.

**Highlights:**

- Irrelevant stimulation to a contralateral body site hinders tactile detection.
- We show robust and early integration of sensory inputs from across body sides.
- The amount of interference varies by the signal-to-noise in the irrelevant stimulation.
- Interference may result from cortical integration of bilateral tactile sensations.

## Introduction

Our environment contains far more information than we can process. Much of that information is irrelevant to our goals, but attention guides our perception by prioritising relevant inputs over distracting interference. While our eyes (and ears) normally work in sync with one another, processing the same inputs simultaneously, our sense of touch does not enjoy the same harmonious coordination. Most of the time, the input to our skin differs substantially across our body, from our feet to our fingers. Sometimes, it must be integrated (for example, when turning the steering wheel of a car with both hands); and sometimes it must be set apart and distinguished (for example, when turning the steering wheel of a car with one hand, while shifting gears with the other). In this sense, the tactile modality is unique from vision and audition, which receive coherent inputs to both eyes or ears. However, there is relatively little research about how simultaneous tactile stimulation to different parts of the body surface is integrated.

The pathways for sensing touch have been extensively delineated in human and in animal models (Y Iwamura 2000; Yoshiaki Iwamura 1998; Vernon B Mountcastle 1957; V B Mountcastle 1997; Mark Tommerdahl, Favorov, and Whitsel 2010), but the integration rules for inputs coming from different parts of the body have been less studied. Physiological studies performed in animal models provide some insight (Tamè et al. 2016; M Tommerdahl, Favorov, and Whitsel 2005; M Tommerdahl et al. 1999), but these rarely link changes in neural activation patterns to perceptual experience or behavioural outcome (i.e., they are not performed in awake, behaving animals). Hence, we are interested in how simultaneous stimulation to different body sites affects the perception of touch; and more specifically, in how irrelevant tactile sensations across the body-midline affect tactile performance in another body site.

To understand how simultaneous stimulation to different body sites affects perceptual experience, it is useful to consider two features of cortical representation that directly affect the integration and differentiation of incoming inputs, both neighbouring and distant: the selectivity and laterality of the somatosensory cortex (Saadon-Grossman, Arzy, and Loewenstein 2020). *Selectivity* indicates how specific a cortical response is to a particular body part, and *laterality* reflects the extent to which the two sides of the body are processed independently. Recent imaging efforts of the somatosensory homunculus reveal several topographic maps which diverge on the parameters of selectivity and laterality (Saadon-Grossman, Arzy, and Loewenstein 2020). For example, BA 3b (an area within S1) is characterised by specialised finger response, but subsequent S1 processing areas (e.g., BA 1 and BA 2) are less selective. There, receptive fields are less finely tuned, and cortical responses are elicited by neighbouring digits, not just by a preferred digit (Yoshiaki Iwamura 1998; Martuzzi et al. 2014; Saadon-Grossman, Arzy, and Loewenstein 2020).

Diminished selectivity between fingers of the same hand is supported not only by physiological studies, but also by behavioural observations. Many studies show that performance on a tactile target is hampered by the addition of other stimulation sites within the same hand (Schweizer et al. 2001; Sherrick 1964; Tamè, Farnè, and Pavani 2011; Tamè, Moles, and Holmes 2014). This basic finding, i.e., a blurred distinction between fingers, is generalised over various tactile stimulation types and tasks.

Across the hands, the degree of perceptual interaction is less researched. Studies point to overlapping neural representations across body sides, suggesting diminished laterality. These are at least partially scaffolded by transcallosal connections between S1 in opposite hemispheres and by bilateral receptive fields in S2 (Y Iwamura 2000), and are further supported by cognitive studies on homologous fingers. Tamè et al. (2012) showed that in somatosensory cortices, neural adaptation measured during a detection task is finger-specific, but indifferent to body-sides, i.e., greater neural adaptation was observed after stimulation to homologous fingers, compared to neighbouring fingers. At a behavioural level, Tamè and colleagues (2011) reported interference in a go/no-go task, where stimulation to the homologous finger was mistaken for stimulation to the target finger (Tamè, Farnè, and Pavani 2011). Similarly, tactile training can transfer from a trained finger to either neighbouring or homologous fingers of the opposite hand (Harrar, Spence, and Makin 2013; Harris, Harris, and Diamond 2001).

The physiological and behavioural synergy between hands begs the question of how tactile stimulation to both sides of the body interacts when (i) presented concurrently and (ii) within the context of ongoing goals. Here, previous studies provide conflicting clues. A wide variety of tasks have shown impairments in performance when contra-lateral stimulation is applied (Braun et al. 2005; Sherrick 1964; Tamè, Farnè, and Pavani 2011; Tamè, Moles, and Holmes 2014; Nguyen et al. 2014), but others show no effects in performance, or even facilitation (Evans and Craig 1991; Craig 1985; Lappin and Foulke 1973). The discrepancies in these findings may be attributed to differences in stimulation type (e.g., whether distractor and target are identical or not; see Nguyen et al. 2014;

Driver and Grossenbacher 1996), body-site (e.g., homologous vs. non-homologus fingers; see Tamè, Farnè, and Pavani 2011; Nguyen et al. 2014), or task-type (Tamè, Moles, and Holmes 2014). In addition, this body of work investigates the fate of stimulus perception under varied degrees of uncertainty, but none of these studies directly measure performance on targets presented amidst ongoing distraction. Stimulation protocols typically include either location uncertainty or presentation in close succession (i.e., masking protocols).

In this study, we investigated whether concurrent tactile stimulation to two body sites can be distinguished or are necessarily integrated. Specifically, we characterised the impact of irrelevant sensory stimulation on tactile detection performed on the dominant index finger. In order to fully describe the dynamics of irrelevant tactile stimulation, we manipulated: (i) the body sites to which the irrelevant stimulation was applied, as well as (ii) the body posture; and in addition, (iii) the intensity of the irrelevant stimulation and (iv) its content (signal vs. noise).

In Experiments 1-3, we characterised and contrasted the influence of irrelevant stimulation to various body sites (contra-lateral index finger, pinky finger, and ankle) on ongoing perception at the dominant index finger. Thus, we were able to quantify the extent of integration between the target finger and a homologous finger, a non-homologous finger, as well as to a an entirely different body part situated in another part of mental and anatomical space. In Experiment 4, we examined the impact of body posture on detection performance: participants received irrelevant stimulation to the homologous finger, with the arm extended and occluded from view. In Experiment 5, we examined how varying levels of (irrelevant) stimulation intensity affect detection performance. In Experiment 6, we examined the impact of receiving relevant (target) stimulation on the (ignored) contra-lateral hand. And last, in a final Experiment 7, we asked whether our observed effects are specific to the target-type that we employed (a decrement in intensity), or whether they generalise to other stimulation types (i.e., an increment in intensity). Across this set of studies, we document robust interference between the different body sites that is not specific to body parts, unaffected by attentional manipulations, and that is extinguished by task-relevant attributes, indicating vast and early pooling of sensory processes in the somatosensory system.

## Methods

The methods for all 7 experiments are described below. In addition, a visual summary of the key elements in the experimental manipulation is provided in Table 1.

**Table 1:**
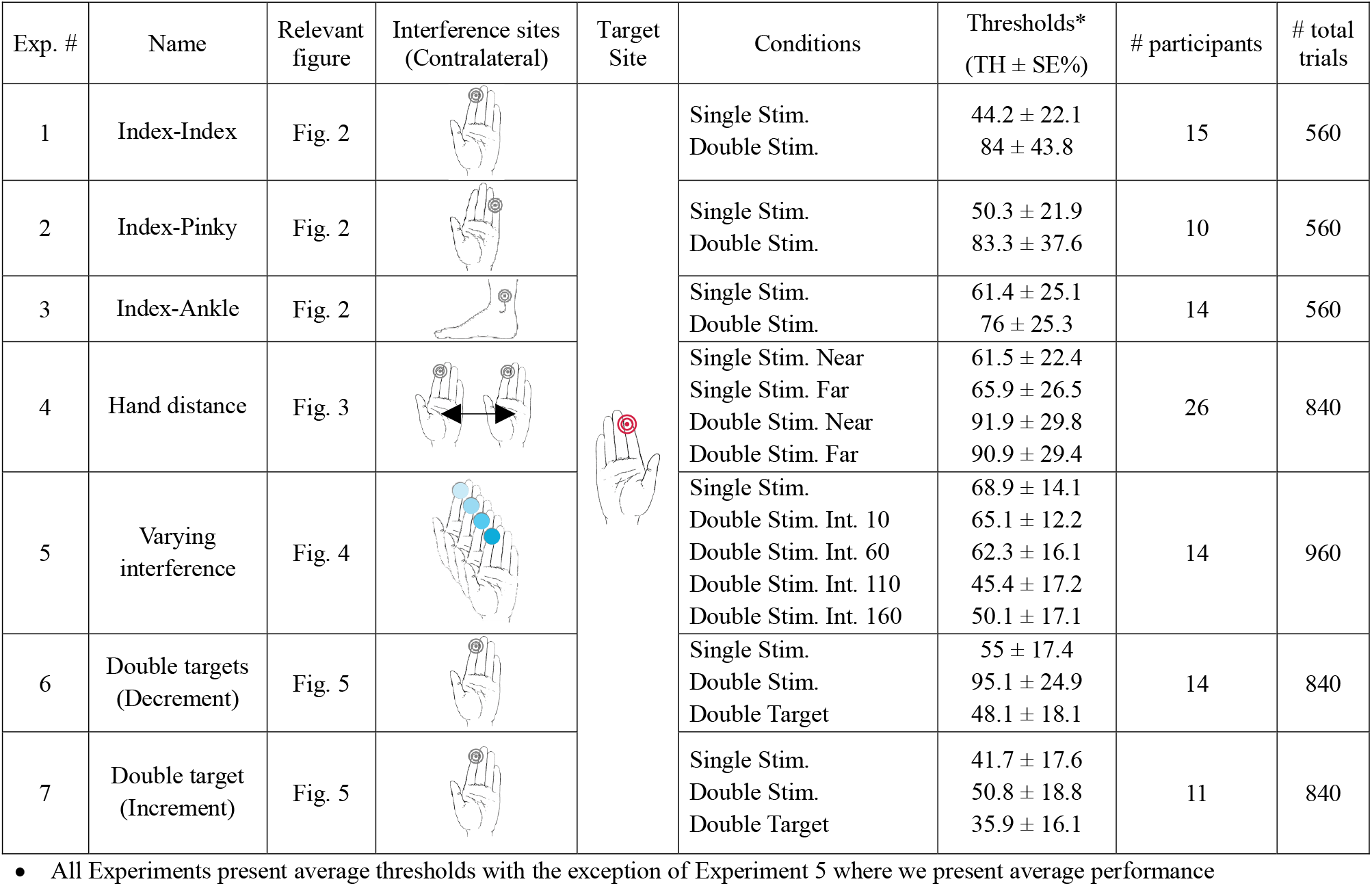
Summary of the experiments

| Exp. # | Name | Relevant figure | Interference sites (Contralateral) | Target Site | Conditions | Thresholds* (TH ± SE%) | # participants | # total trials |
| --- | --- | --- | --- | --- | --- | --- | --- | --- |
| 1 | Index-Index | Fig. 2 | | | Single Stim.<br>Double Stim. | 44.2 ± 22.1<br>84 ± 43.8 | 15 | 560 |
| 2 | Index-Pinky | Fig. 2 | | | Single Stim.<br>Double Stim. | 50.3 ± 21.9<br>83.3 ± 37.6 | 10 | 560 |
| 3 | Index-Ankle | Fig. 2 | | | Single Stim.<br>Double Stim. | 61.4 ± 25.1<br>76 ± 25.3 | 14 | 560 |
| 4 | Hand distance | Fig. 3 | | | Single Stim. Near<br>Single Stim. Far<br>Double Stim. Near<br>Double Stim. Far | 61.5 ± 22.4<br>65.9 ± 26.5<br>91.9 ± 29.8<br>90.9 ± 29.4 | 26 | 840 |
| 5 | Varying interference | Fig. 4 | | | Single Stim.<br>Double Stim. Int. 10<br>Double Stim. Int. 60<br>Double Stim. Int. 110<br>Double Stim. Int. 160 | 68.9 ± 14.1<br>65.1 ± 12.2<br>62.3 ± 16.1<br>45.4 ± 17.2<br>50.1 ± 17.1 | 14 | 960 |
| 6 | Double targets (Decrement) | Fig. 5 | | | Single Stim.<br>Double Stim.<br>Double Target | 55 ± 17.4<br>95.1 ± 24.9<br>48.1 ± 18.1 | 14 | 840 |
| 7 | Double target (Increment) | Fig. 5 | | | Single Stim.<br>Double Stim.<br>Double Target | 41.7 ± 17.6<br>50.8 ± 18.8<br>35.9 ± 16.1 | 11 | 840 |

- All Experiments present average thresholds with the exception of Experiment 5 where we present average performance

### Experiment 1: Homologous Finger

#### Participants

We pre-specified a sample size of 20 participants based on previous psychophysical and tactile interference studies. Twenty-one healthy participants took part in a psychophysical experiment for payment or class credit. Two participants were excluded due to a technical problem. One participant was excluded due to poor performance. Three others were excluded because the target hand turned out to be their non-dominant hand. Data are thus presented for 15 participants (ages 19-26, average = 23; 11 females; all right-handed). Participants signed a consent form before experimentation and the study was approved by the Institutional Review Board of Human Experimentation at The Hebrew University of Jerusalem.

#### Apparatus

Stimulation was produced with a vibrotactile coin stimulator connected to an open-source hardware, Arduino (Uno Rev3), programmed with C++ on compatible IDE. The experiment was built and run on OpenSesame v3.1 (Mathôt, Schreij, and Theeuwes 2012). The vibration produced by the Arduino was approximately 120 Hz. Headphones were used to administer white noise throughout the experiment, in order to prevent participants from hearing the vibration. Data analyses were conducted using Matlab 2017b (The MathWorks, Inc. Natick, MA, USA) and the Palamedes toolbox (Prins and Kingdom 2018). This apparatus served all reported experiments.

#### Stimuli

Stimulation consisted of an ongoing constant vibration that lasted 1.6 s. The detection target was embedded within the ongoing stimulation and consisted of a brief (0.04 s) decrement in intensity (Figure 1a). We use arbitrary units (AU) to describe the intensity of the vibration and ΔAU to describe the decrement, where Δ = constant intensity - target intensity. The intensity of the ongoing constant vibration was 160 AU and that of the decrements (i.e., the targets) were parametrically varied, ranging from Δ20 to Δ140 AU, in steps of 20. This resulted in 7 target intensity levels: Δ20, Δ40, Δ60, Δ80, Δ100, Δ120, Δ140, from difficult to easily detectable targets, respectively. In certain cases, two additional intensities were used (Δ10 and Δ150), but as all participants did not have these extreme values, they were excluded from analyses. Target onset was randomised from 0.5 to 1.1 s within the 1.6 s long stimulation following the onset of the vibrotactile stimulation.

**Figure 1.**
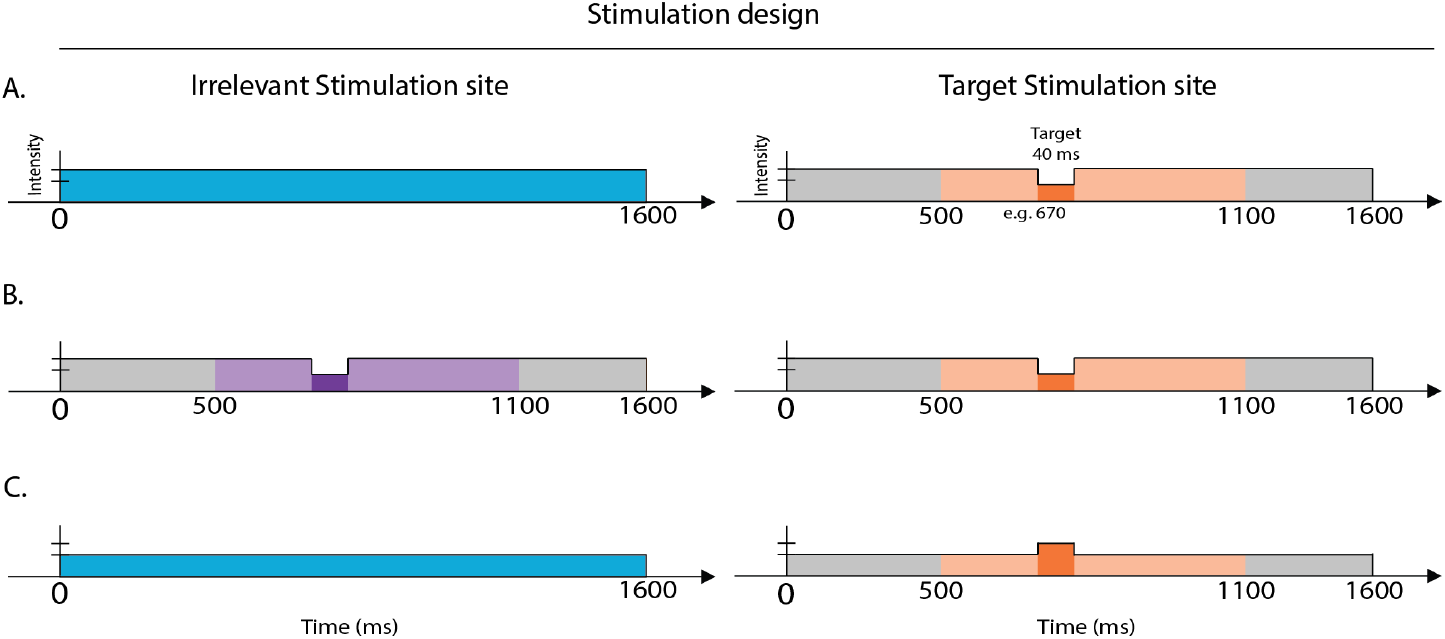
Stimulation Design. In all experiments, participants received ongoing (1.6 s) vibrotactile stimulation to the index finger of their dominant hand (target stimulation site) and to an additional, contralateral body site (irrelevant stimulation site). The target, a 0.04-s change in the intensity, was embedded in half of the trials (50% catch trials). In target present trials, detection targets appeared at a random time point between 0.5 and 1.1 s after vibration onset. The intensity change varied parametrically, resulting in seven target levels ranging from hardly-to easily-detectable. After the end of the 1.6-s stimulation, participants were prompted to indicate via foot pedals whether or not they had detected the target. In Experiments 1-6, the target was an intensity *decrement* (panels a and b), while in Experiment 7 the target was an intensity *increment* (panel c). In Experiments 6 and 7, the irrelevant stimulation site contained an intensity change equivalent to the target in half of all target-present trials (panel b). Dark orange (or purple) denote the brief decrement in stimulus intensity. Light orange (or purple) denotes the time range within which targets could appear, by design.

#### Task

Participants were instructed to focus on a fixation cross while attending to their dominant hand (the stimulation site). In each trial, an ongoing vibrotactile stimulation was delivered to participants’ dominant index finger, at the end of which a response screen prompted them to indicate whether they had felt the embedded target or not via foot pedals. They were instructed that there would be easy and hard trials, as well as trials without a target at all (catch trials).

Participants pressed the right foot pedal for “yes” (using their right foot) and the left foot pedal for “no” (using their left foot). To keep false alarm rates to a minimum, we instructed participants to be conservative: to only answer “yes” when they were “quite sure” that they had felt the target, and to otherwise answer “no.”

Trials in which the vibrotactile stimulation was administered only to the dominant (target) index finger are referred to as Single Stimulation trials. Trials in which vibrotactile stimulation was delivered to both the dominant (target) index finger, as well as to an additional (irrelevant) site (here, the homologous finger) are referred to as Double Stimulation, or Irrelevant Simultaneous Stimulation (ISS) trials. The additional site, unless otherwise noted, always vibrated at the intensity of the ongoing vibration (160 AU), and for the entire duration of trial (1.6 sec). It was identical to the vibration administered to the target finger, but without an embedded target (unless otherwise noted).

Single and Irrelevant Simultaneous Stimulation (ISS) conditions were balanced (equal number of trials) but were randomised within target intensity blocks (i.e., target intensities were blocked). Each intensity block included 40 trials with a target, and 40 trials with no target (i.e., 50% catch trials per target intensity). This led to a total of 560-720 trials. The order of catch and target-present trials was randomized. The blocked target intensities were presented in randomized order, except the first, which was always the easiest (i.e., Δ140, or the second easiest, for those few participants who also performed Δ150; see Procedure).

#### Procedure

After signing an informed consent form, participants were seated in front of a computer monitor, 75 cm from the screen, with both arms positioned on the chair’s armrest. Participants’ wrists were supported but the hands themselves made no contact with the armrest (i.e., were hanging off the armrest). A vibrotactile stimulator was secured to their dominant index fingertip with a customized, elastic bandage (target hand). A second vibrotactile stimulator was attached to the body site designated as the interference site (in this Experiment, the contralateral index finger) and this vibration coin vibrated with the same temporal properties as the target vibration coin, only without the embedded target. The computer monitor showed a white cross centrally positioned against a black background that served as a fixation cross throughout the experiment, and the response options appeared on either side of it, corresponding to their positions on the foot pedals (i.e., “no” on the left side of the cross and “yes” on the right).

After instructions were given, participants underwent a practice phase, consisting of 32 trials. Here, participants were familiarised with the ongoing, constant stimulation, as well as with target-present trials. Then, they completed a short practice block (32 total trials: 8 trials for each of the two easiest intensities, in which half were One-Hand and half were SS trials; and 16 catch trials). Practice was repeated if participants scored either <80% hit rate or >20% false alarm rate. All participants met this criterion after a maximum of three repetitions. The experimenter remained present in the room during the practice phase.

After completing the practice, participants wore headphones and listened to white noise while they performed the experiment. All participants began with an easy target intensity (Δ140), after which performance was assessed. If they met the criterion (as in the training, i.e., ≥80% hit rate and ≤20% false alarm rate), they continued on to the rest of the experiment. Otherwise, participants repeated this block in order to ensure that they understood the task. If they still failed to meet the criterion, they were given an even easier block (Δ150 target intensity), after which performance was re-assessed. If participants still failed to meet the criterion, they were excused from the experiment. Otherwise, they continued on to the rest of the experimental blocks, which were presented in randomised order.

The interval between trials was set to 1 s, with a 0.25 s jitter (1-1.25 s). There was a short optional break every 80 trials, which was prompted by a screen indicating that participants could take a short break and press any foot pedal to continue, when ready. At the end of the experiment, participants were either paid for their time or granted class credit.

### Experiment 2: Non-Homologous Finger

#### Participants

We pre-specified a sample size of 10 participants, indicated by a power analysis based on Experiment 1. Ten healthy participants took part in a psychophysical experiment for payment or class credit. No participants were excluded. Data are presented for 10 participants (ages 19-38 average = 24; 8 females; 7 right-handed). Participants signed a consent form before experimentation and the study was approved by the Institutional Review Board of Human Experimentation at The Hebrew University of Jerusalem.

#### Irrelevant Stimulation Site

All experimental parameters were identical to Experiment 1, with the exception of the additional (irrelevant) stimulation site, which was the pinky finger of the non-dominant hand.

### Experiment 3: Contra-lateral Ankle

#### Participants

We pre-specified a sample size of 15 participants, again based on a power analysis derived from the previous Experiments, 1 and 2. Fifteen healthy participants took part in a psychophysical experiment for payment or class credit. One participant was excluded due to a technical problem. Data are presented for 14 participants (ages 20-28, average = 23; 10 females; 12 right-handed). Participants signed a consent form before experimentation and the study was approved by the Institutional Review Board of Human Experimentation at The Hebrew University of Jerusalem.

#### Irrelevant Stimulation Site

All experimental parameters were identical to Experiments 1 and 2, with the exception of the additional (irrelevant) stimulation site, which was the ankle contralateral to the target stimulation site (i.e., the dominant hand).

### Experiment 4: Variable Body Posture

#### Participants

We pre-specified a sample size of 30 participants based on previous psychophysical and tactile interference studies. Thirty-three healthy participants took part in a psychophysical experiment for payment or class credit. Seven participants were excluded; one due to a technical problem, two did not complete the experiment, and four due to poor performance (below chance). Data are thus presented for 26 participants (ages 19-35, average = 24; 15 females; 21 right-handed). Participants signed a consent form before experimentation and the study was approved by the Institutional Review Board of Human Experimentation at The Hebrew University of Jerusalem.

#### Stimulus

Identical to Experiments 1-3, with the exception that here, no extra (easier or more difficult) intensities were used, resulting in the seven target intensities specified in Experiment 1 (Δ20 to Δ140 AU, in steps of 20).

#### Task

Identical to Experiments 1-3, with the exception of (i) an additional body posture manipulation (i.e., in which the hands were set farther apart, with the non-target hand extended and occluded from view) and (ii) number of repetitions. With respect to (i), the experiment was divided into two main blocks: in one, the hands were positioned close together, as in Experiments 1 and 2; and in the other block, the hands were set farther apart, with the non-target arm extended and occluded from view. The order of these two blocks (Hands-Close; Hands-Far) was counter-balanced across participants. In each block, participants were presented all 7 target intensities (blocked, as in Experiments 1-3). Within each target intensity block, the ratio of Single to Double Stimulation trials to catch trials was 1:2:3 (i.e., 50% catch trials, and twice the Double Stimulation trials as Single Stimulation trials). This resulted in 30 trials with no target and 30 trials with a target (20 Double and 10 Single Stimulation trials). This led to a total of 420 trials per Body Posture (Hands-Close, Hands-Far), or a total of 840 trials in the entire experiment. As in Experiments 1-3, the order of catch and target-present trials was randomised, as was the order of target intensities.

#### Procedure

Identical to Experiments 1-3, with the exception of the practice phase, which again consisted of 32 total trials, but was split into two parts. In the first half, participants’ arms were placed as in previous experiments. In the second half, the non-target arm was extended and occluded from view. It rested on a small platform just below shoulder height, with hands and fingers hanging off the platform. All other parameters remained the same, including the frequency of breaks.

### Experiment 5: Variable Irrelevant Stimulation Intensity

#### Participants

We pre-specified a sample size of 20 participants based on previous psychophysical and tactile interference studies. Twenty participants took part in a psychophysical experiment for payment or class credit. Six participants were excluded; one did not complete the experiment, three due to poor performance (below chance) across intensity levels; and two due to extremely high false alarms in at least one condition (69% and 100%). Data are thus presented for 14 participants (ages 18-26, average = 22; 10 females; 13 righthanded). Participants signed a consent form before experimentation and the study was approved by the Institutional Review Board of Human Experimentation at The Hebrew University of Jerusalem.

#### Stimulus

Identical to Experiments 1-4, with the exception that here, only four target intensities were used (40-100 ΔAU, in steps of 20). In addition, the intensity of the additional (irrelevant) stimulation site (i.e., non-target hand) varied, such that there were four levels of irrelevant stimulation intensity: 160 AU, as in the previous experiments; and 110, 60, and 10 AU. Including the Single Stimulation condition, this resulted in five Stimulation conditions: one Single Stimulation condition, and four Double Stimulation conditions.

#### Task

Similar to Experiments 1-4, but here the Stimulation conditions were blocked (i.e., the Single and the four Double Stimulation conditions). The Single Stimulation condition was always presented first, followed by the Double Stimulation condition with the highest stimulation intensity (i.e., 160 AU, the same Double Stimulation condition as was used in the previous experiments). The other three Double Stimulation conditions were presented in randomised order. Within each Stimulation block, participants experienced the full range of target intensities (blocked and presented in randomised order). There were 20 repetitions per target intensity, and an equal number of catch trials (50% catch trials). This led to a total of 160 trials per Stimulation Intensity block, or 800 total trials in the experiment.

#### Procedure

Similar to the previous experiments, with the exception of the practice block. In this experiment, participants were again familiarised with the ongoing, constant stimulation, as well as with target-present trials. However, then they completed a short practice block in which 4 total trials were presented in the Single Stimulation condition only (2 trials for each of the two easiest intensities, and 2 catch trials). Then, they completed another short practice block in which 8 total trials were presented in the Double Stimulation condition only (4 trials with the easiest two intensities and with the simultaneous (irrelevant) stimulation at the intensity of the ongoing constant vibration, i.e., 160 AU; and 4 catch trials). All other parameters were identical to previous experiments.

### Experiment 6: Double Target

#### Participants

We pre-specified a sample size of 20 participants based on previous psychophysical and tactile interference studies. Twenty-one participants took part in a psychophysical experiment for payment or class credit. Three participants were excluded did not complete the experiment, and four others were excluded due to poor performance (below chance) or ceiling performance across intensity levels. Data are thus presented for 14 participants (ages 20-28, average = 23; 9 females; 11 right-handed). Participants signed a consent form before experimentation and the study was approved by the Institutional Review Board of Human Experimentation at The Hebrew University of Jerusalem.

#### Stimulus

Identical to the previous experiements, except that a third experimental condition was added, in which an intensity decrement identical to the target was embedded in the simultaneous (irrelevant) stimulation (i.e., the non-target finger). In this experimental condition, the intensity decrement always had the same intensity and temporal characteristics as the target presented to the dominant hand (i.e., co-occurred with the target). This experimental condition (Double Target) occurred in addition to the Single and Double Stimulation conditions. (See Figure 1b)

#### Task

Similar to previous experiments. Single, Double Stimulation, and Double Target conditions were balanced (equal number of trials) and randomised within target intensity blocks. Each intensity block included 60 trials with a target (i.e., 20 repetitions per Stimulation condition), and 60 trials with no target (i.e., 50% catch trials). This led to a total of 840 trials.

#### Procedure

Identical to previous experiments. The practice block was identical to that in Experiment 1.

### Experiment 7: Double Target, Target-Increment

#### Participants

We pre-specified a sample size of 10 participants based on previous psychophysical and tactile interference studies. Fourteen participants took part in a psychophysical experiment for payment or class credit. Three participants were excluded due to ceiling or floor performance, or lack of a dynamic range of performance across intensity levels. Data are thus presented for 11 participants (ages 19-27, average = 23; 5 females; 9 righthanded). Participants signed a consent form before experimentation and the study was approved by the Institutional Review Board of Human Experimentation at The Hebrew University of Jerusalem.

#### Stimulus

Identical to Experiment 6 (Double Target), except that, in this experiment, the detection target was a brief (0.04 s) *increment* in intensity (rather than a decrement in intensity). The intensity of the ongoing constant vibration was 100 AU and that of the increments (i.e., the targets) were parametrically varied, ranging from Δ20 to Δ140 AU, in steps of 20. This resulted in 7 target intensity levels: Δ120, Δ140, Δ160, Δ180, Δ200, Δ220, Δ240, from difficult to easily detectable targets, respectively. (See Figure 1c)

#### Task

Identical to Experiment 6.

#### Procedure

Identical to previous experiments. The practice block was identical to that in Experiments 1 and 6.

### General

#### Analyses

For each participant included in the analysis, we estimated the percentage of hits, misses, correct rejections and false alarms for each condition (e.g., Single and Double Stimulation conditions) separately.

For all main analyses, we used a two-tailed percentile bootstrap procedure for dependent groups, with 10,000 samples with replacement (Efron and Tibshirani 1993; Wilcox 2012). We compared mean detection rates between Stimulation conditions (e.g., Single vs. Double Stimulation; target intensities, collapsed) by calculating percentile confidence intervals around the mean difference between the two stimulation-conditions. First, we sampled participants with replacement, keeping their corresponding mean detection rates (i.e., averaged across target intensity levels) in the two conditions. We then calculated the mean difference between conditions across all (sampled) participants. We performed these two steps 10,000 times, and each time saved the mean difference between conditions. Then, we sorted the bootstrapped means, and used the 2.5 and 97.5 percentiles to form the boundaries of the 95% bootstrap confidence intervals (for an alpha, α = 0.05). To calculate whether the detection rates in the two conditions of interest differed from each other, we estimated the overlap of the bootstrapped distribution with zero (i.e., the null hypothesis, that there is no difference in detection rates between the two stimulation-conditions), in the following manner: p-value = [one minus the percentage of bootstrap values above (or below) zero, multiplied by two (for a two-tailed test)].

To estimate psychometric functions, the responses (hit rates, or percent detection) for each participant were modelled by fitting logistic functions for each experiment, using a maximum-likelihood procedure for each condition (e.g., Single, Double Stimulation) and experiment (Palamedes toolbox; Prins and Kingdom 2018). We then calculated the 50% threshold of performance, as well as the slope of the psychometric curve, for each condition separately. Both threshold (α) and slope (β) parameters were allowed to vary freely, while guess (γ=0) and lapse (λ=0.01) rates were fixed for all participants.

A post-hoc analysis between Experiments 1-3 contrasted the magnitude of interference between pairs of Experiments, in order to determine whether the impact of the irrelevant stimulation varied as a function of stimulation site (i.e., homologous index finger, non-homologous pinky finger, and contralateral ankle). The magnitude of interference in each experiment is given by the difference in 50%-thresholds between the Single and Double Stimulation conditions in that experiment. We then used a two-tailed percentile bootstrap procedure for independent groups to contrast the difference in magnitude between all possible pairs (three). First, we sampled participants with replacement, using the minimum number of participants across the three experiments (n = 10, Experiment 2). We then calculated the mean difference in 50%-thresholds across all sampled participants (for each Experiment separately), and computed the difference between Experiments. We performed these steps 10,000 times, and each time saved the difference between Experiments. Then, we sorted the bootstrapped differences, and used the 2.5 and 97.5 percentiles to form the boundaries of the 95% bootstrap confidence intervals (as outlined earlier in this section). To calculate whether the interference magnitudes in the two experiments of interest differed from each other, we estimated the overlap of the bootstrapped distribution with zero (i.e., the null hypothesis, that there is no difference in magnitude of interference between the two experiments), in the following manner: p-value = [one minus the percentage of bootstrap values above (or below) zero, multiplied by two (for a two-tailed test)].

In Experiment 5 (Varying ISS Intensity), we modelled the relationship between the ISS intensity and target detection performance using a simple linear regression. This analysis was performed for each target intensity separately. Thus, for a given target intensity we fitted a first degree polynomial to detection performance over the different ISS intensity levels (see Figure 4, panel b). This was performed for each participant separately and yielded four slopes per participant (one for each target intensity). Then, we resampled participants with replacement, and derived a sampling distribution of average-slope values for each target intensity (with 2.5 and 97.5 percentiles to form the boundaries of the 95% bootstrap confidence intervals).

To calculate whether these slopes differed from those that would be observed if ISS intensity did not significantly impact target detection performance, we computed a bootstrap sampling distribution based on shuffling ISS intensity labels and generating slopes based on the shuffled data. This surrogate analysis was also performed within each target intensity level separately.

First, we shuffled ISS intensity labels for each participant, and computed the slope using the linear regression method described above. We repeated this procedure 10,000 times and averaged the slopes across participants for each iteration. This results in a distribution of group averaged slopes. This bootstrap sampling distribution was contrasted with the distribution of the original slope data, by computing the overlap.

## Results

### Experiment 1: Homologous Finger

Figure 2a (top left panel) shows the mean psychometric functions in the Single and Double (homologous index finger) stimulation-conditions. As per experimental design, participants’ performance increased as a function of increasing target intensities, from nearly no target detection at the hardest intensity levels to almost perfect target detection at the easiest intensity levels. To examine the effects of irrelevant stimulation on target detection, we first compared overall mean detection rates between the two stimulation-conditions (Single, Double); and subsequently the 50%-thresholds of each.

**Figure 2.**
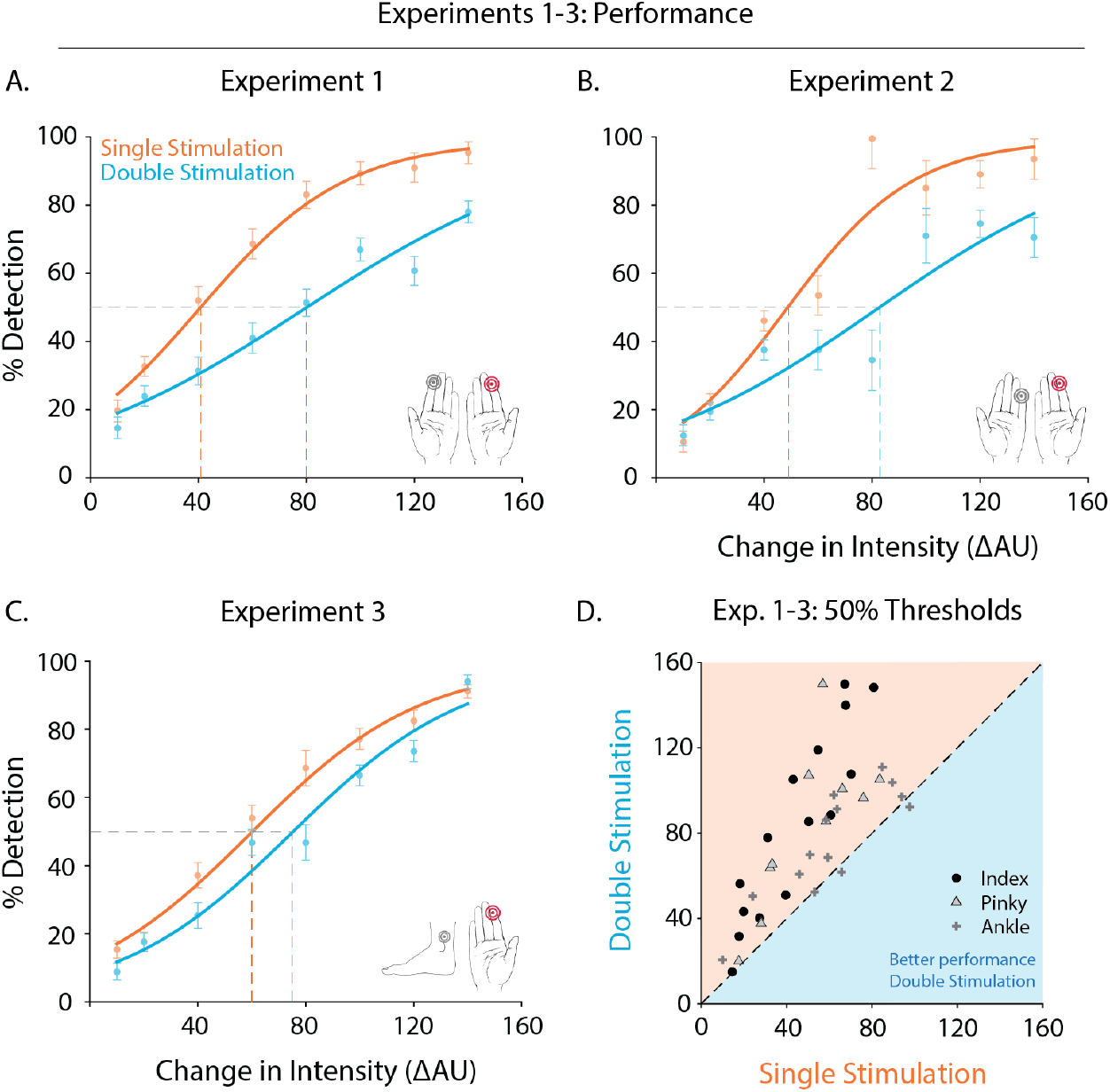
Detection Performance for Experiments 1 – 3. Mean psychometric functions in the Single and Double Stimulation conditions for Experiments 1 (panel a), 2 (panel b), and 3 (panel c). In all three experiments, participants’ ability to detect tactile targets while receiving additional (irrelevant) stimulation to a contralateral body site was significantly hindered. The scatterplot (panel d) shows the 50% thresholds for individual participants (single points) in all three experiments. Nearly all participants exhibited lower thresholds in the Single stimulation-condition (almost all participants to the left of the identity line).

For all main analyses, we used a two-tailed percentile bootstrap for dependent groups, with 10,000 samples with replacement. We compared overall mean detection rates between the two stimulation-conditions (Single vs. Double Stimulation; Target Intensity, collapsed) by calculating percentile confidence intervals around their difference (see Methods; General; Analyses). Square brackets indicate the boundaries of the 95% confidence intervals (CI) constructed from this analysis.

In Experiment 1, participants generally exhibited higher detection rates when targets were presented to the dominant index finger with no additional simultaneous stimulation, compared to when they received irrelevant simultaneous stimulation to the homologous finger (Single minus Double Stimulation = 22.7% [16.7, 28.6], p < .001).

In addition, we compared target intensities corresponding to the 50% detection rate in each stimulation-condition, as estimated from individual participants’ psychometric fits. Here too, participants exhibited lower detection thresholds in the Single Stimulation condition (mean 50%-threshold, Single minus Double Stimulation = - 39.8 ΔAU [−52.0 - 27.6], p < .001). This difference corresponds to that of two target-intensity levels higher in the Double Stimulation condition. All participants showed this pattern (see Figure 2d).

In addition, the difference in performance between stimulation-conditions also manifested as a difference in slopes between individual participants’ psychometric fits, such that the slope of the Single stimulation-condition was consistently higher than that of the Double stimulation-condition (mean slopes, Single minus Double Stimulation: 0.027 [.008 .051], p < .005). There was no difference in false alarms between stimulation-conditions (mean false alarm rate across the entire experiment, 13.8 % ± 9.8, p = 0.4).

### Experiment 2: Non-Homologous Finger

In Experiment 2, we conducted the same Experiment as above, but placed the additional vibrotactile stimulator on a non-homologous finger: the contralateral pinky finger. Not only is this the farthest possible finger from the dominant index finger, but it is also served by a different dermatome. In order to rule out interference effects resulting from sensory overlap in the spinal tract (e.g., due to decussation), we opted to stimulate a digit connected to an entirely different spinal nerve.

Figure 2b (top right panel) shows the mean psychometric functions in the Single and Double (contralateral pinky finger) stimulation-conditions. As per experimental design, participants’ performance again improved as a function of increasing target intensities. Like in Experiment 1, we compared overall mean detection rates between the two stimulation-conditions, and subsequently the 50%-thresholds of each, in order to examine the effects of the irrelevant stimulation on target detection.

Participants again exhibited higher detection rates when targets were presented to the dominant index finger without additional stimulation, compared to when they received irrelevant simultaneous stimulation to the contralateral pinky finger (Single minus Double Stimulation = 19.9% [14.5, 26.2], p < .001).

Participants also exhibited lower detection thresholds in the Single compared to the Double stimulation-condition (mean 50%-threshold, Single minus Double Stimulation = - 33.0 ΔAU [−49.9, −19.8], p < .001). This difference corresponds to that of one-to-two target-intensity levels higher in the Double Stimulation condition. All participants showed this pattern (see Figure 2d).

The difference in performance between stimulation-conditions also manifested as a difference in slopes between individual participants’ psychometric fits, such that the slope of the Single Stimulation condition was consistently higher than that of the Double Stimulation condition (mean slopes, Single minus Double Stimulation: 0.038 [.023, .056], p < .001). There was no difference in false alarms between stimulation-conditions (mean false alarm rate across the entire experiment, 8.3 % ± 6.1, p = 0.6).

### Experiment 3: Contralateral Ankle

Experiment 3 was identical to Experiments 1 and 2, but here the irrelevant simultaneous stimulation was administered to the contralateral ankle.

Figure 2c (bottom left panel) shows the mean psychometric functions in the Single and Double (contralateral ankle) stimulation-conditions. As per experimental design, participants’ performance again improved as a function of increasing target intensities. Like in Experiments 1 and 2, participants exhibited higher detection rates when targets were presented to the dominant index finger without additional stimulation, compared to when they received simultaneous stimulation to the contralateral ankle (Single minus Double Stimulation = 7.9% [4.2, 11.6], p < .001). This difference in detection-performance also manifested as lower detection thresholds for the Single stimulation-condition (mean 50%-threshold, Single minus Double Stimulation = −14.6 ΔAU [−21.2 - 8.1], p < .001). This difference corresponds to that of almost one target-intensity level higher in the Double stimulation-condition. Nearly all participants showed this pattern (see Figure 2d).

In contrast to Experiments 1 and 2, there was no difference in slopes or false alarms between stimulation-conditions (mean slope across the entire experiment, 0.04 ± 0.02, p = 0.4; mean false alarm rate across the entire experiment, 8.2 % ± 6.5; p = 0.2).

### Experiment 4: Variable Body Posture

In this experiment, participants underwent the same protocol as in Experiments 1 (i.e., the irrelevant stimulation site was the homologous index finger), but during half of the experiment, the irrelevant stimulation site (i.e., the non-target arm) was extended and occluded from view.

Figure 3 shows the mean psychometric functions in both Single and Double stimulation-conditions, in Near and Far arm postures (i.e., both hands in front of the body, as in Experiments 1-3; or non-target arm extended and behind the body, occluded from view). As per experimental design, participants’ performance increased as a function of increasing target intensities, from nearly no target detection at the hardest intensity levels to almost perfect target detection at the easiest intensity levels.

**Figure 3.**
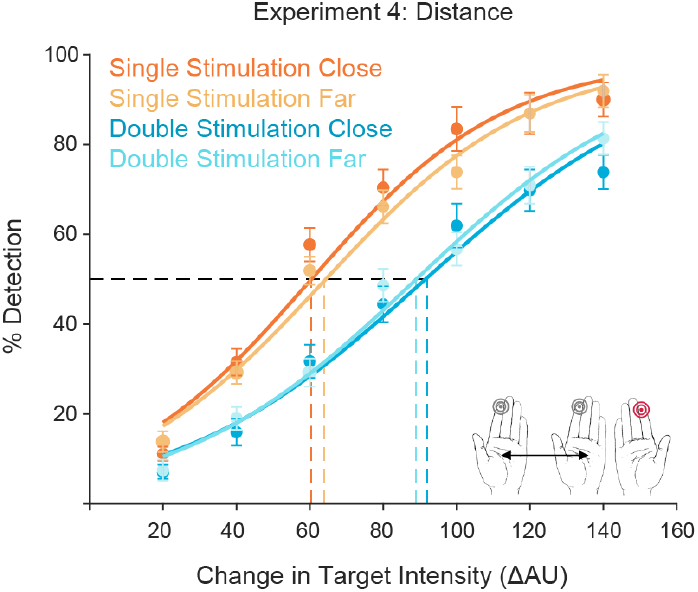
Detection Performance for Experiment 4. Mean psychometric functions in the Single and Double Stimulation conditions for Experiment 4, in Near and Far arm postures (dark orange and dark blue, or light orange and light blue respectively). In both body postures, participants’ ability to detect tactile targets while receiving additional stimulation to the homologous finger was significantly and equally hindered.

We assessed whether the position of the hands (Near, Far) impacts the magnitude of detection-interference. To this end, we first compared overall mean detection rates (target intensity, collapsed) between the two stimulation-conditions when the arms were positioned far apart (Single vs. Double Stimulation, Hands-Far). We observed a robust impact on target-detection, as observed in all previous experiments (Single minus Double Stimulation, Hands-Far = 14.3 % [11.3, 17.9], p < 0.001). In addition, and as observed in Experiments 1-3, we observed a robust impact on target-detection when the hands were positioned close together (Single minus Double Stimulation, Hands-Near, 18.0 % [15.2, 20.9], p < 0.001).

Within stimulation-conditions, the overall mean detection rates between Far and Near body postures did not differ (Double Stimulation: Hands-Far minus Hands-Near, p = 0.6; Single Stimulation: Hands-Far minus Hands-Near, p = 0.3).

We then compared target intensities corresponding to the 50% detection rate, as estimated from individual participants’ psychometric fits. Participants exhibited lower detection thresholds in the Single compared to the Double stimulation-condition, in both body postures (50%-thresholds; Single minus Double Stimulation, Hands-Far: −25.1 ΔAU [−31.5 −19.2], p < .001; Hands-Near: −30.4 ΔAU [−35.7 −25.2], p < .001). As observed in Experiments 1-3, this difference in 50%-thresholds corresponds to that of one-to-two target-intensity levels higher in the Double stimulation-condition. Within stimulation-conditions, the 50%-thresholds did not differ between the two body postures (Double Stimulation, p = 0.8; Single Stimulation, p = 0.2).

The above differences in detection-performance also manifested as a difference in slopes between individual participants’ psychometric fits, such that the slope of the Single stimulation-condition was consistently higher than that of the Double stimulation-condition (mean slopes; Single minus Double Stimulation, Hands-Far: 0.013 [.0043 .0234], p = .003; Hands-Near: 0.018 [0.0072, 0.0297], p = .001). Within stimulation-conditions, the slopes did not differ between body postures (Double Stimulation, p = 0.6; Single Stimulation, p = 0.4).

Unlike in previous experiments, participants exhibited different false alarm rates between stimulation-conditions, such that there were more false alarms in the Single compared to the Double stimulation-condition (mean false alarm rate across the entire experiment, 8.23 % ± 7.9; Single minus Double Stimulation, Hands-Far: 3.02 % [1.51, 4.67], p < .001; Hands-Near: 4.26 % [2.55, 6.21], p < 0.001). Within stimulation-conditions, there was no difference in false alarms between body postures (Double Stimulation, p = 0.7; Single Stimulation, p = 0.2).

### Interim Summary

In Experiments 1-4, we examined the impact of irrelevant tactile stimulation on tactile detection in a constantly attended location. We found a compelling effect of interfering stimulation on detection performance for all body sites that we measured (homologous and non-homologous fingers, and contra-lateral ankle).

In order to assess whether the effects of detection-interference differed as a function of the additional (irrelevant) stimulation site, we contrasted the magnitude of interference between experiments 1-3. The magnitude of interference is given by the difference in 50%-thresholds between stimulation-conditions. This analysis indicated that the impact of interference between Experiments 1 (homologous finger) and 2 (non-homologous finger) did not differ (mean 50%-thresholds; Experiment 2 minus Experiment 1 = 6.78 ΔAU [−14.5, 27.3], p = 0.5). However, the magnitude of interference when the additional stimulation was applied to the contralateral ankle (Experiment 3) was significantly less than that of Experiments 1 and 2 (mean 50%-thresholds; Experiment 3 minus Experiment 1 = 25.2 ΔAU [8.2, 42.6], p < .003; Experiment 3 minus Experiment 2 = 18.4 ΔAU [8.3, 42.3], p = .003).

In summary, the impact of irrelevant tactile stimulation on detection-performance was not modulated by finger identity or finger distance from the target site, but it was alleviated by applying the additional stimulation to the contralateral ankle – in the order of one target intensity level lower. The impact of irrelevant tactile stimulation on detection-performance was not modulated by body posture: even when the non-target hand was fully extended and occluded from view, detection-performance did not change as compared to when the hand was in front of the body and within view.

### Experiment 5: Variable Irrelevant Stimulation Intensity

In this experiment, participants underwent a similar protocol as in Experiment 1 (i.e., the irrelevant stimulation site was the homologous index finger). However, participants performed the experiment with only four target intensities, and with an additional experimental manipulation in which the irrelevant simultaneous stimulation (ISS; i.e., to the non-target hand) varied in intensity. Thus, the non-target hand could vibrate at one of four different intensity levels (10, 60, 110, or 160 AU). We examined the impact of the Double Stimulation intensity on detection-performance.

Figure 4 (panel a) shows the mean performance across target intensity levels for the five stimulation-conditions (Single Stimulation; and the four Double Stimulation intensity levels described above). As per experimental design, participants’ performance increased as a function of increasing target intensities, from very little or below-chance target detection at the hardest intensity levels to greater target detection at the easiest intensity levels.

**Figure 4.**
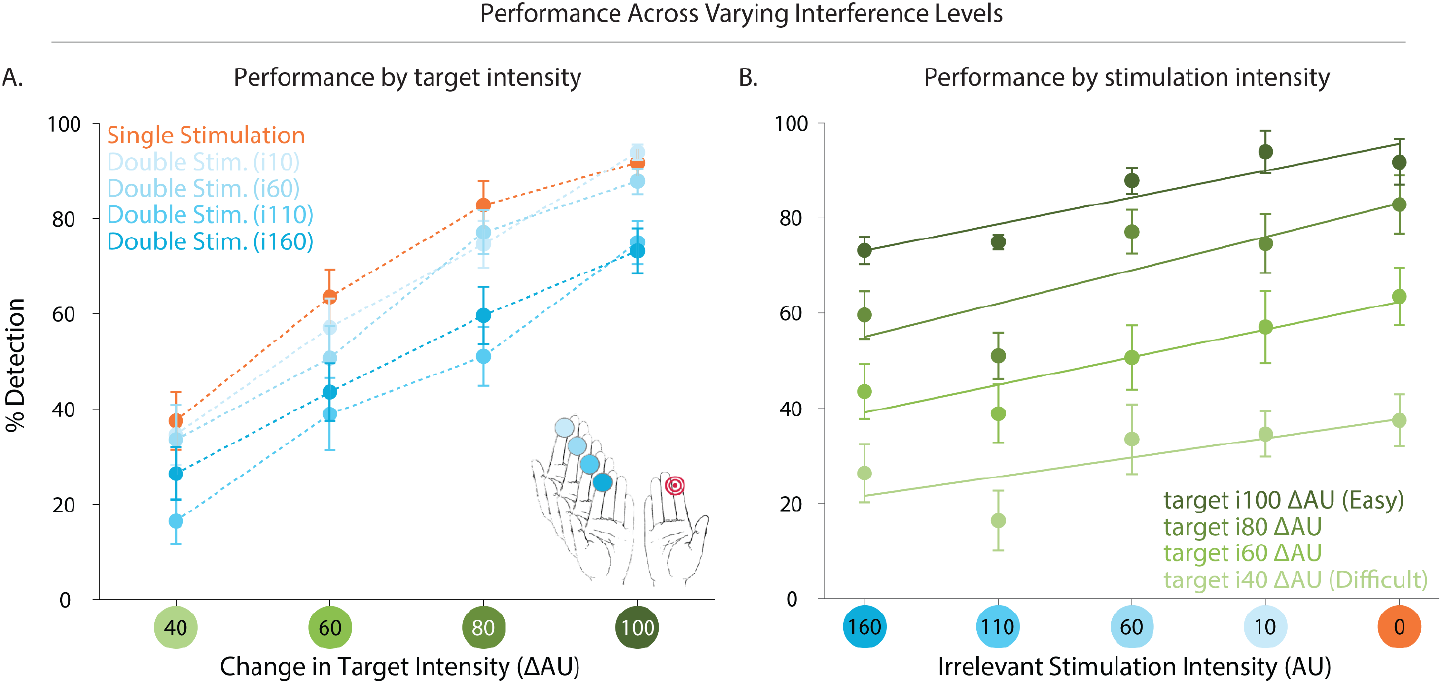
Detection Performance in Experiment 5. Participants’ ability to detect tactile targets varied as a function of target intensity (panel a) and irrelevant stimulation intensity (panel b). Participants received different levels of irrelevant stimulation intensity to the homologous index finger. For each irrelevant stimulation intensity, participants exhibited higher detection rates as target intensity increased (panel a). The relationship between detection-performance and irrelevant stimulation intensity can be modelled as a simple linear regression, for each target intensity level (panel b), such that lower irrelevant stimulation led to increased detection-performance, except for in the weakest target intensity level (panel b, lightest green line).

In order to estimate the impact of irrelevant stimulation intensity on detection-performance, we modelled the relationship between the two using a simple linear regression (see Figure 4, panel b). This analysis was performed for each target intensity level, separately. A comparison between the derived slopes and a bootstrap null distribution of slopes (see Methods; General; Analyses) indicated that detection-performance increased linearly with decreased irrelevant stimulation intensity, except for in the weakest target intensity (see Figure 4, panel b, lightest green line) (mean slopes across participants: 100 ΔAU = 0.14 [.11, .21], p < .001; 80 ΔAU = 0.16 [.11, .21], p < .001; 60 ΔAU = 0.13 [.063, .20], p < .003; 40 ΔAU, p = 0.6).

### Experiment 6: Double Target

In this Experiment, we added a third stimulation-condition, referred to as the Double Target condition. In this condition, an intensity decrement of the same magnitude as the target and with the same temporal characteristics was embedded in the irrelevant stimulation (see Figure 1b).

Figure 5a shows the mean psychometric functions in the Single, Double, and Double Target stimulation-conditions. As per experimental design, participants’ performance again increased as a function of increasing target intensities, from almost no target detection at the hardest intensity levels and progressively increasing until reaching nearly perfect target detection at the easiest intensity levels. To examine the effects of the Double Target on detection performance, we first compared overall mean detection rates between the three conditions; and subsequently the 50%-thresholds of each.

**Figure 5.**
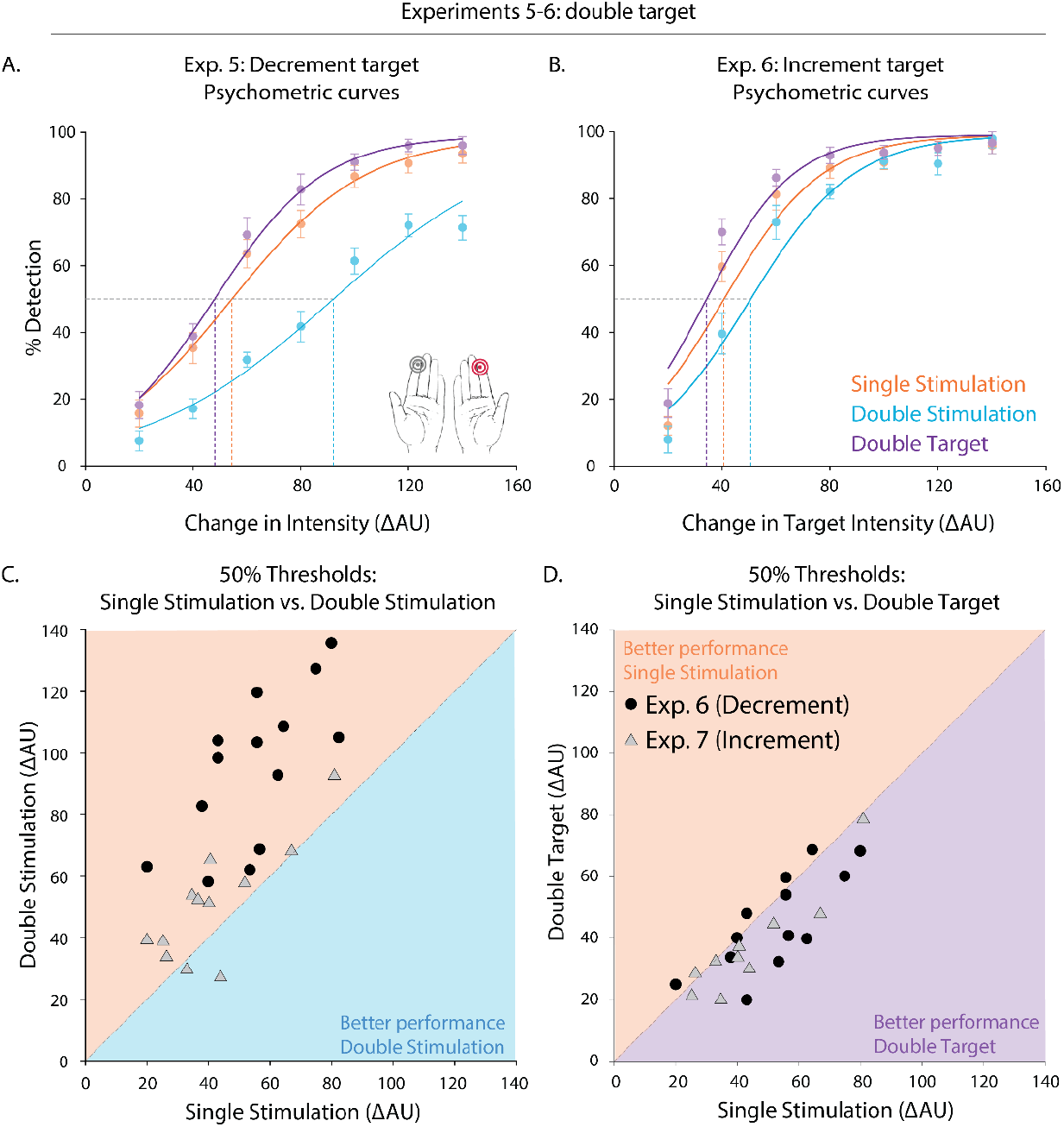
Detection Performance for Experiments 5 and 6. Mean psychometric functions in the three stimulation-conditions for Experiments 5 (panel a) and 6 (panel b). In both experiments, participants’ ability to detect tactile targets while receiving additional (irrelevant) stimulation to the homologous index finger was significantly hindered. The scatterplots (panel c and d) shows the 50% thresholds for individual participants (single points) in the Single and Double stimulation-conditions (panel c); and in the Single and Double Target stimulation conditions (panel d). Nearly all participants exhibited lower thresholds in the Single, as compared to the Double stimulation-condition (panel c; almost all participants to the left of the identity line); but even lower thresholds in the Double Target stimulation-condition (panel d; most participants to the right of, or along, the identity line).

In order to assess the impact of the Double Target on detection-performance, we compared overall mean detection rates between the Double Target and the Single and Double stimulation-conditions (target intensity, collapsed). We observed a significant difference in detection-performance in the Double Target as compared to both Single and Double stimulation-conditions, such that participants exhibited higher detection rates in the Double Target condition (Double Target minus Single Stimulation: 4.9 % [1.8 8.2], p < .005; Double Target minus Double Stimulation: 27.0 % [22.8, 31.0], p < .001). As expected, participants also exhibited higher detection rates in the Single compared to the Double stimulation-condition (Single minus Double Stimulation: 22.1 % [17.1, 27.2], p < .001). We also compared target intensities corresponding to the 50% detection rate in each interference condition, as estimated from individual participants’ psychometric fits. Here too, participants exhibited lower detection thresholds in the Double Target condition relative to the others (Single Stimulation minus Double Target: 6.9 ΔAU [1.6, 12.5], p < .009; Double Stimulation minus Double Target: −46.9 ΔAU [−56.2, −38.0], p < .001). Additionally, and as expected, participants exhibited lower detection thresholds in the Single as compared to the Double Stimulation (Single minus Double Stimulation: - 40.1 ΔAU [−48.8 −30.3], p < .001). While nearly all participants exhibited their lowest detection rates in the Double stimulation-condition, most of them also exhibited their highest detection rates in the Double Target condition (see Figure 5, panels c and d).

These differences in detection-performance also manifested as a difference in slopes between individual participants’ psychometric fits. The slope of the Single Stimulation was higher than that of the Double stimulation-condition (mean slopes, Single minus Double Stimulation: 0.017 [.0082 .027], p < .001); but lower than that of the Double Target condition (Single minus Double Target Stimulation: −.023 [−.045, −.0027], p < .03). As expected, the slope of the Double Target was also higher than that of the Double Stimulation condition (Double Target minus Double Stimulation: .040 [.022, .063], p < .001). There was no difference in false alarms (mean false alarm rate across the entire experiment, 1.8 % ± 2.2) between any of the stimulation-conditions (Single vs. Double Stimulation, p = 0.9; Single vs. Double Target Stimulation, p = 0.8; Double Target vs. Double Stimulation, p = 0.5).

### Experiment 7: Intensity Increment – Double Target

This Experiment is identical to the previous one, except that here the target was always an increment in intensity, rather than a decrement (see Figure 1c).

Figure 5b shows the mean psychometric functions in the Single, Double, and Double Target Stimulation conditions. As per experimental design, participants’ performance increased as a function of increasing target intensities. To examine the effects of the Double Target condition on detection-performance, we again compared overall mean detection rates between the three conditions; and subsequently the 50%-thresholds of each.

The findings of this experiment mirrored those of Experiment 6. Participants showed hindered detection-performance in the Double stimulation-condition (Single minus Double Stimulation: 6.0 % [2.9, 9.2], p < .001), and enhanced performance in the Double Target condition (Single minus Double Target conditions, −4.2 % [−6.7, −2.0], p < .001).

Here too, participants exhibited lower detection thresholds in the Double Target condition relative to the others (Single Stimulation minus Double Target: 5.8 ΔAU [2.4, 9.7], p < .001; Double Stimulation minus Double Target: −14.9 ΔAU [−20.8, −8.9], p < .001). Again and as expected, participants exhibited lower detection thresholds in the Single Stimulation compared to the Double Stimulation (Single minus Double Stimulation: −9.1 ΔAU [−14.9 −2.5], p < .009). While nearly all participants exhibited their lowest detection rates in the Double Stimulation condition, many of them also exhibited their highest detection rates in the Double Target condition (see Figure 5, panels c and d).

The differences in detection performance between the three stimulation-conditions manifested as a difference in slopes between individual participants’ psychometric fits in the Single Stimulation versus Double Target comparison (Single minus Double Target Stimulation: −.026 [−.041, −.0099], p < .001), but not in any of the other stimulation-condition comparisons (Single minus Double Stimulation, p = 0.6; Double Target minus Double Stimulation, p = 0.1). There was no difference in false alarms (mean false alarm rate across the entire experiment, 1.8 % *±* 2.1 between any of the stimulation-conditions (Single vs. Double Stimulation, p = 0.6; Double Target vs. Double Stimulation, p = 0.3), except for between the Single and Double Target stimulation-conditions (0.71 % [0.12, 1.4], p < .03)

## Discussion

In this set of studies, we investigated how simultaneous stimulation to two sites on opposite sides of the body impact tactile cognition. To this end, we ran a series of experiments in which individuals had to detect a target on their dominant index finger, while receiving irrelevant stimulation to another body site. We asked whether concurrent tactile inputs – one always relevant and the other always irrelevant – can be differentiated in accordance with ongoing goals, or whether they are necessarily integrated. Understanding how multiple tactile inputs interact informs our ecological reality: in our daily life, the tactile receptors across our skin surface, from our feet to our hands, are constantly stimulated by dissimilar objects, and depending on our current goals, they must sometimes be coordinated, and sometimes set apart.

Previous tactile studies investigating the impact of stimulation across the hands focused primarily on short-duration stimuli (in the order of .01-.2 sec). Here, we aimed to more closely mimic the longer-duration tactile conditions encountered in everyday life. Thus, we applied ongoing (1.6 sec) vibration to fingers of opposite hands, one of which contained a brief (0.04 sec) embedded intensity change (the target). Participants were instructed to ignore the irrelevant stimulation site and to detect the target, thus enabling an examination of contralateral stimulation with complete certainty over target location. We found that participants’ ability to detect tactile targets was impaired, regardless of whether distractor vibration was applied to the homologous or non-homologous effector, or even to a different limb. In contrast to previous studies involving a wide variety of tasks (e.g., discrimination, localisation, or go/no-go protocols), our participants performed a simple detection task. Our choice of task enabled a more direct examination of how well a tactile target is detected in light of irrelevant stimulation.

### A Role for Attention versus Integration in Tactile Interference

In our first set of studies, we applied irrelevant stimulation to the homologous finger (non-dominant index finger; Exp 1), and subsequently to the contralateral pinky (Exp 2) and contralateral ankle (Exp 3) while participants performed a tactile detection task. In all cases, participants’ ability to detect the target was impaired, even though they were explicitly instructed to focus only on the target-hand and to ignore the irrelevant stimulation site because the target would never occur there. In all three experiments, all participants clearly displayed reduced detection rates and higher detection thresholds compared to single-site stimulation, meaning that the irrelevant stimulation interfered with their ability to detect the target (see Figure 2). The observed interference was significantly greater in magnitude when produced by the contralateral hand (homologous or non-homologous finger) compared to the contralateral ankle, which is more distant both in physical space and in the neural representation.

Experiment 4 (Variable Body Posture) sought to address the origin of these observed interference effects. One possibility is that they result from an involuntary attentional capture by the body site receiving irrelevant stimulation (i.e., it cannot be ignored), thus eliciting sensory competition between the two body sites (Johansen-Berg and Lloyd 2000). Alternatively, stimulation from the two body sides might be integrated early along the pathway of somatosensory processing (i.e., reduced laterality in the cortical response) and lead to a degraded capacity to differentiate concurrent stimulations.

Thus, in Experiment 4, participants’ non-target arm (i.e., receiving irrelevant stimulation) was extended and occluded from view, minimising the potential role of involuntary attentional capture. Attentional shifts between body parts have been shown to take time in tactile detection tasks (Lakatos and Shepard 1997). In addition, numerous studies have reported that non-informative vision of a body part (i.e., in this experiment, the target arm) improves tactile detection thresholds and discrimination performed on that body part, potentially by affecting tactile receptive fields (Haggard et al. n.d.; Kennett, Taylor-Clarke, and Haggard 2001; Schaefer, Heinze, and Rotte 2005). Thus, if attentional capture were the main driver of the interference effects, we would expect for occlusion of the irrelevant arm to result in attentional facilitation and increased sensitivity in the target-hand, leading to enhanced performance when the hands were further apart. However, in our experiment individuals’ tactile performance was unaffected by the distance between hands. Even with the contralateral arm extended and occluded from view (i.e., was further away and in a different part of space than the target hand), the detrimental effects of irrelevant stimulation remained unchanged. These results suggest that the observed tactile interference cannot be attributed solely to attentional effects. Instead, the tactile input to both hands may be integrated along the somatosensory cortical pathways, leading to diminished laterality in tactile processing.

### How is tactile stimulation across contra-lateral body sites combined/ integrated?

Experiments 5 and 6 allowed us to understand detection performance under different combinations of signal (target hand) and noise (irrelevant hand). In Experiment 5, we parametrically varied the intensity of the irrelevant stimulation, such that in some experimental blocks it was less intense (i.e., lighter) and in others more intense (i.e., closer to the ongoing intensity of the target hand). As the intensity of the irrelevant stimulation increased, individuals exhibited worse detection performance (see Figure 4). In fact, higher levels of noise masked the target-signal in a linearly increasing manner.

Experiment 6 complemented these results by giving us insight into how concurrent tactile stimulation interacts when the irrelevant body site unpredictably contains relevant target information (i.e., a concurrent intensity change of equal magnitude as the target). A simple summation between stimulation from the two body sites would predict that no changes in detection performance should be observed (i.e., double the signal, and double the noise in the Double Target condition, resulting in the same 1:1 signal-to-noise ratio as in the Single Stimulation condition). However, as shown in Figure 5, the presence of a double target (i.e., an equal intensity change both in relevant and irrelevant body sites) enhanced detection performance such that detection was consistently and significantly better than in the Single Stimulation condition. This suggests that integration between concurrent stimuli may be more complex than a simple summation. Electrophysiological studies support this notion, as amplitudes following concurrent stimulation are smaller than the sum of each individual stimulation (Severens et al. 2010), and strong concurrent stimuli result in a suppressive interaction between fingers (Gandevia, Burke, and Mckeon 1983).

### Generalising beyond a Stimulus-Type (Decrement Change)

In our last experiment (7), participants underwent the same task as described above (Experiment 6), but in which the target was an intensity increment (rather than a decrement) relative to the ongoing vibration. We observed the same effects as in Experiment 6, allowing us to generalise our findings beyond the target stimulus (i.e., a decrement). Interestingly, the magnitude of the interference effects was inferior to that observed with a target decrement. This observation is in line with phenomenological reports that detecting an increment is easier than detecting a decrement. It is also possible that the SNR ratios between the noise and the target increment, and between the noise and the target decrement, were not phenomenologically equivalent between our experiments. Nonetheless, presence tasks are generally easier than absence tasks (Whang, Burton, and Shulman 1991), also as predicted by Weber’s law.

### Conclusion

In this set of tactile studies, we set out to understand how concurrent stimulation to opposite body sites affects tactile cognition. In most of our daily life, we are bombarded by simultaneous tactile inputs across our skin surface, the signals of which must be differentiated, coordinated or ignored by our somatosensory system in accordance with our ongoing goals. Across the seven experiments reported here, we characterise robust tactile interference when receiving irrelevant stimulation to the opposite body side during detection performance on a known, attended body part. This interference is consistently present across individuals and is not modulated by attentional manipulations (e.g., changes in body posture), pointing to an early pooling of tactile inputs that results in an integrated percept. A recent neuroimaging study describes reduced laterality in the cortical response of the somatosensory system (Saadon-Grossman, Arzy, and Loewenstein 2020). Our work supplements this neural response characterization with a thorough description of its functional consequences to detection in somatosensation. Consistent with reduced laterality, we demonstrate robust and early integration of sensory inputs from across body sides. The rules for integration across body sides are likely not described by a simple summation, but rather may be governed by more complex interactions between fingers and according to the corresponding perceived as well as actual intensities of the stimulation. Future studies combining time resolved neuroimaging with sophisticated psychophysics might shed light on the precise locus and rules of integration in the somatosensory system.

## Acknowledgments

This work was supported by funds from the James McDonnell Scholar Award for Understanding Human Cognition (awarded to A.N.L.) and the Jerusalem Brain Community (awarded to S.P.). The authors would like to thank Gal Moscona for her theoretical and practical contributions to this work; as well as Lorenzo Pisoni and Niv Pagir for their technical contributions to the experimental setup; and Carolyn Saito, Deena Nerwen, Meira Robbins, Daniel Tomer, Elad Stephenson, Sapir Suissa and Daniel Aped for their help with experimental setup and data collection.

